# Structural Mechanisms of Actin Isoforms

**DOI:** 10.1101/2022.08.01.502282

**Authors:** Amandeep S. Arora, Hsiang-Ling Huang, Ramanpreet Singh, Yoshie Narui, Andrejus Suchenko, Tomoyuki Hatano, Sarah M. Heissler, Mohan K. Balasubramanian, Krishna Chinthalapudi

## Abstract

Actin isoforms organize into distinct networks that are essential for the normal function of eukaryotic cells. Despite a high level of sequence and structure conservation, subtle changes in their design principles determine the interaction with myosin motors and actin-binding proteins. The functional diversity is further increased by posttranslational modifications (PTMs). Therefore, identifying how the structure of actin isoforms relates to function is important for our understanding of normal cytoskeletal physiology. Here, we report the high-resolution structures of filamentous skeletal α-actin (3.37Å), cardiac α-actin (3.07Å), ß-actin (2.99Å), and γ-actin (3.38Å) in the Mg^2+^·ADP state with their native PTMs. The structures revealed isoform-specific conformations of the N-terminus that shifts closer to the filament surface upon myosin binding, thereby establishing isoform-specific interfaces. Retropropagated structural changes further show that myosin binding modulates actin filament structure. Further, our structures enabled us to reveal the location of disease-causing mutations and to analyze them with respect to known locations of PTMs. Collectively, the previously unknown structures of single-isotype, posttranslationally modified bare cardiac α-actin, ß-actin, and γ-actin reveal general principles, similarities, and differences between isoforms. They complement the repertoire of known actin structures and allow for a comprehensive understanding of *in vitro* and *in vivo* functions of actin isoforms.

## Introduction

Actin isoforms are among the most ubiquitous and abundant structural proteins that facilitate the functional organization of the cytoplasm of eukaryotic cells (1, 2). Humans express six actin genes in a tissue-specific and developmentally-regulated manner (3). The gene products are structurally and functionally conserved among eukaryotes and can be grouped into four muscle actins (skeletal α-actin, smooth α-actin (vascular), and cardiac α-actin, smooth γ-actin (enteric)) and two nonmuscle actins (ß-actin and γ-actin) (4, 5). Actin isoforms display differential biochemistries, cellular localization, and interactions with myosin motors and actin-binding proteins (ABPs) (2, 3, 6–10). These characteristics result in the formation of diverse cellular actin networks with distinct compositions, architectures, dynamics, and mechanics that enable fundamental processes such as cell adhesion, migration, and contractility (1, 11, 12). Actin isoforms share a high sequence identity at the protein level (∼93-99%) and the ability to self-assemble into helical, polarized filaments (F-actin) from monomers (G-actin) (2, 13–15). Since the publication of the first crystal structure of G-actin in complex with DNaseI ∼ 30 years ago, extensive studies have advanced our understanding of the structure of monomeric and filamentous actin, polymerization mechanisms, posttranslational modifications (PTMs), interaction with drugs, myosin motors, and ABPs at ever-increasing resolution (14, 16–31). Despite these important advances, isoform-specific mechanisms with myosin motors and ABPs that drive cellular functions and biochemistries are widely documented but poorly understood.

To address how the structure contributes to the functional distinction of actin isoforms, we employed recombinant and native protein production approaches to obtain pure, single-isotype preparations of individual isoforms to perform cryo-electron microscopy (cryo-EM) analyses. The 2.99-3.38 Å-resolution structures of filamentous actin isoforms with their native PTMs confirmed the overall design principles, and revealed similarities as well as differences between them. We show that the defining feature used to regulate the interaction with binding proteins is the divergent N-terminus that shows isoform-specific conformations that result in distinct binding regions for myosin motors and ABPs. By comparing bare with previous myosin-bound actin structures, we found that once the actin-myosin interface is formed, the N-terminus is shifted closer to the filament surface, thereby establishing isoform-specific binding interfaces. Next, we employed comparative structural analysis to dissect whether the binding of myosin motors alters actin filament structure in an isoform-dependent manner. We found that myosin motors induce subtle conformational changes in the actin filament with pronounced changes in the N-terminus and DNaseI loop (D-loop) that are likely to contribute to different biochemical properties and distinct actin landscapes in the cell. Lastly, our high-resolution structures of native actin isoforms allowed us to precisely map PTMs and human disease-causing mutations to establish connections between genetic variation and protein function. Together, our work will inform biochemical and cell biological discoveries of isoform-specific actin regulation mechanisms.

## Results

### High-resolution structures and general principles of actin isoforms

To determine the structural characteristics and differences between actin isoforms, we solved the high-resolution structures of single-isotype skeletal α-actin (3.37 Å), cardiac α-actin (3.07 Å), ß-actin (2.99 Å), and γ-actin (3.38 Å) in the filamentous state using cryo-EM (***Figure 1A-D, Figure 1-figure supplement 1, Table 1***). The structures show local resolutions ranging from 2.1-3.5 Å for skeletal α-actin, 1.9-3.3 Å for cardiac α-actin, 1.7-3.0 Å for ß-actin, and 2.1-4.4 Å for γ-actin (***Figure 1-figure supplement 2***). For all actin isoforms, the highest local resolutions were obtained in the central core region. Lower local resolutions were obtained in the most surface exposed and flexible regions such as the N-terminus and the D-loop as expected. All actin structures were solved in the Mg^2+^·ADP state (Figure 1E, Figure 2) and have been obtained without the use of stabilizing drugs that may interfere with the filament structure (6, 32, 33).

**Figure 1:**
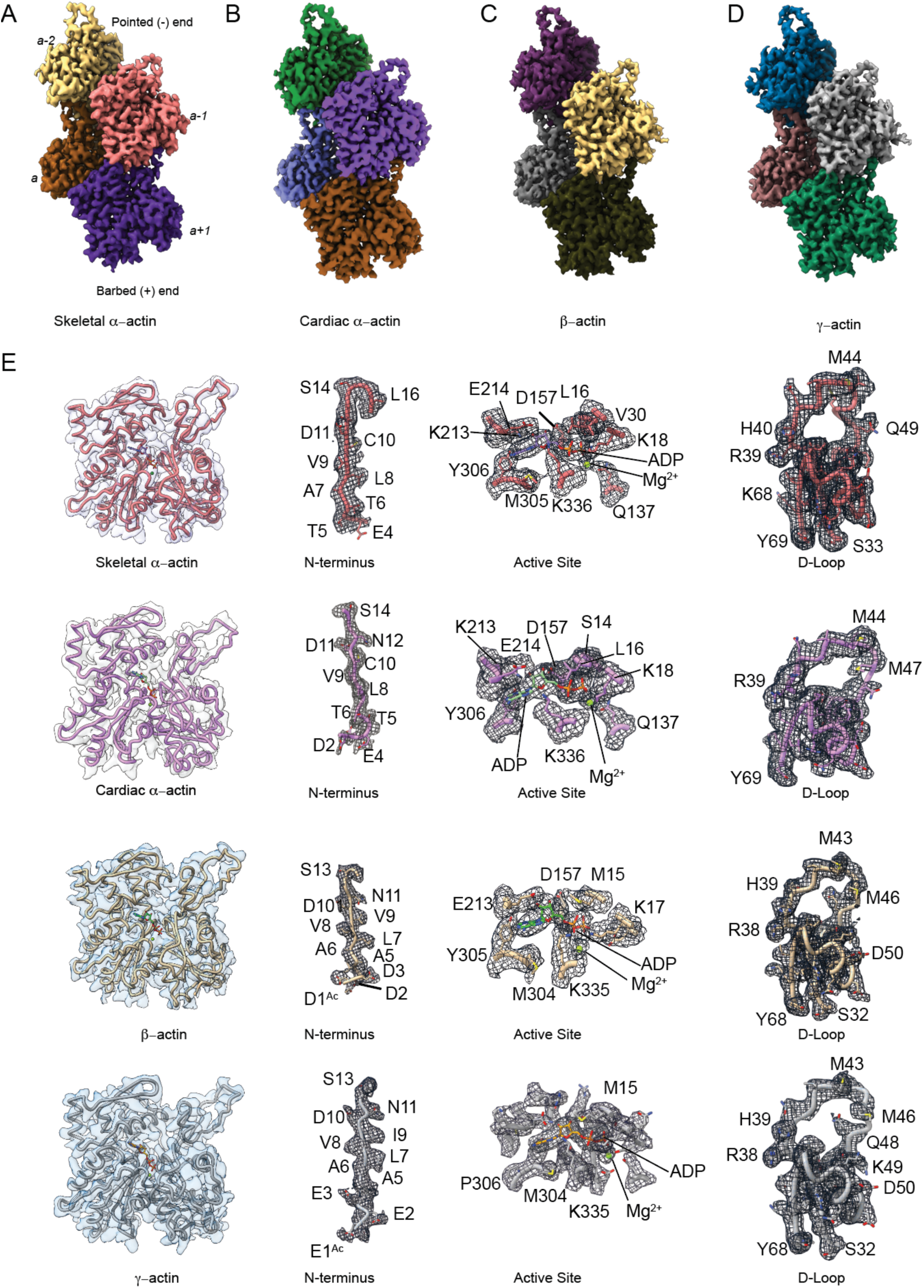
Cryo-EM filament structures of actin isoforms. (A) Helical reconstruction of skeletal α-actin, (B) cardiac α-actin, (C) β-actin, (D) and γ-actin. Views in (B)-(D) are according to (A). Four individual actin protomers in the filament are shown and denoted with italic numbers. The pointed (-) and barbed (+) ends are indicated. (E) Representative cryo-EM densities of key regions of actin isoforms are displayed. Throughout this work, amino acids are numbered according to the sequence of mature actin isoforms (***Figure 1-figure supplement 1C***).

**Figure 2:**
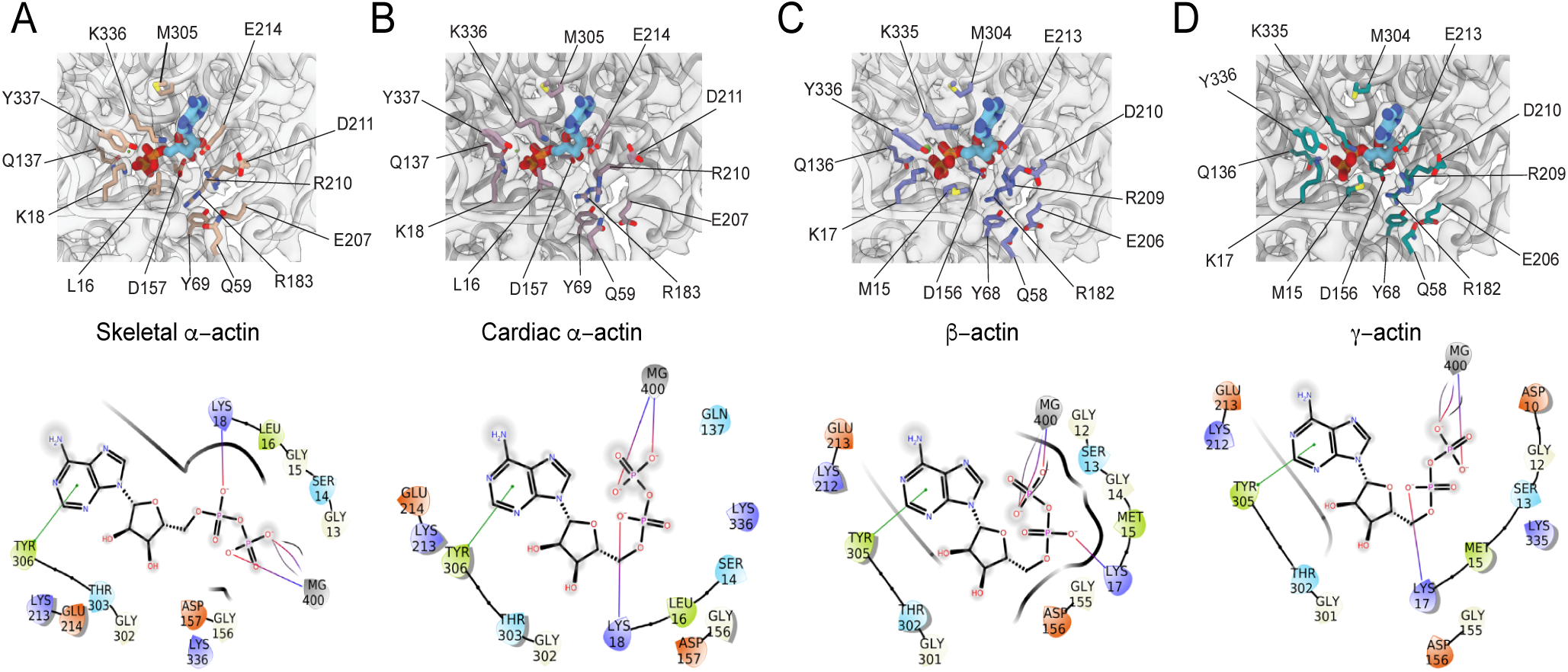
Conserved active site in the actin isoforms. (A) Coordination of Mg^2+^·ADP in the nucleotide-binding cleft of skeletal α-actin. (B) Coordination of Mg^2+^·ADP in the nucleotide-binding cleft of cardiac α-actin. (C) Coordination of Mg^2+^·ADP in the nucleotide-binding cleft of β-actin. (D) Coordination of Mg^2+^·ADP in the nucleotide-binding cleft of γ-actin. ADP is shown in cyan-colored stick representation. Electron densities for key amino acids in the active site of actin isoforms are shown. Schematic representations of interactions in the active sites of the respective actin isoforms are shown in the lower panel. Positively charged amino acids are shown in blue, negatively charged amino acids are shown in red, hydrophobic amino acids are shown in green, polar amino acids are shown in cyan, and Mg^2+^ is shown in gray color. The schematics are not drawn to scale.

**Table 1:**
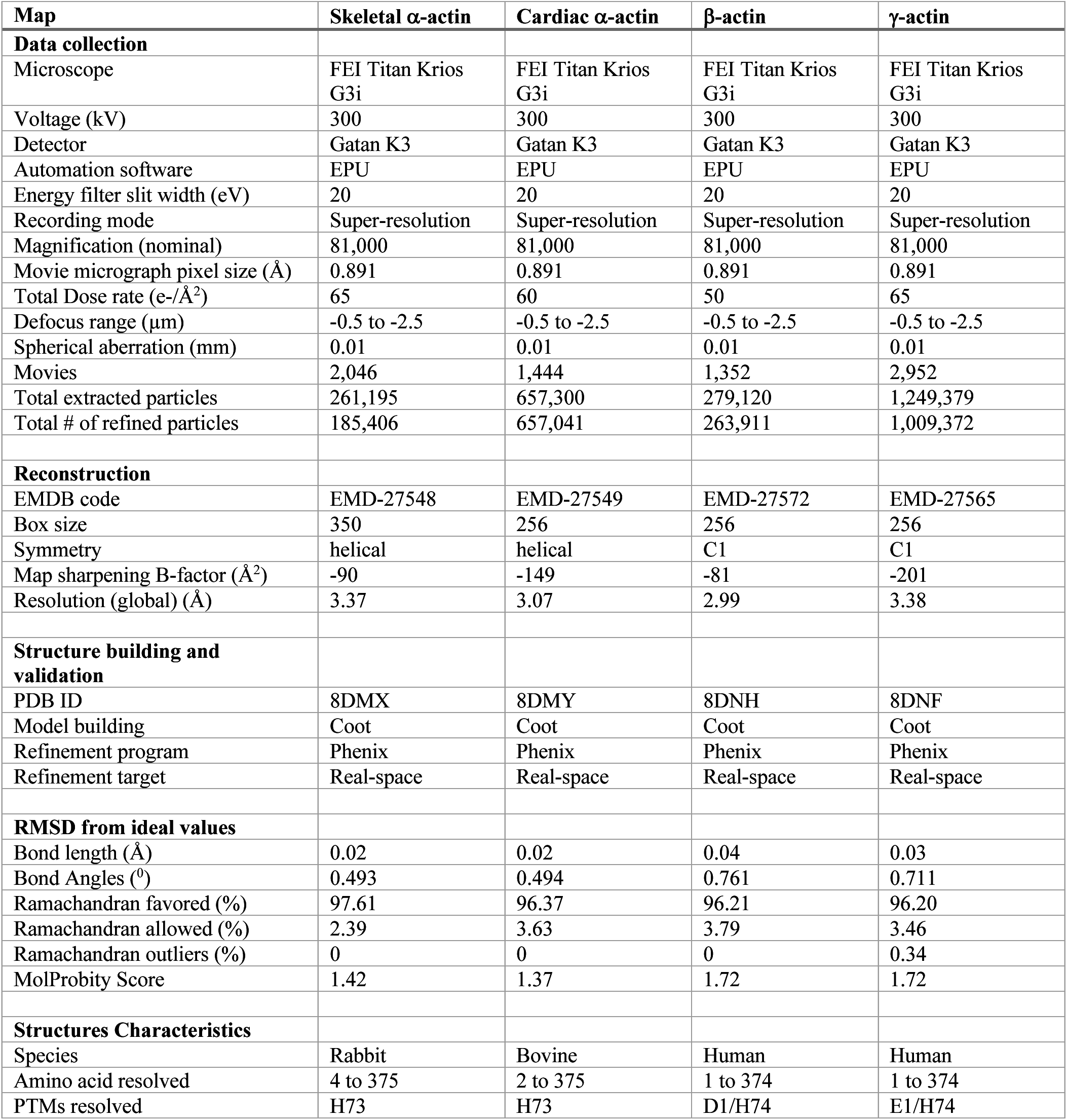
Data collection, image processing, and structure characteristics summary.

Our cryo-EM maps allowed us to build unambiguous models of actin isoforms in which secondary structure information including the side chains, the nucleotide and associated cation (Mg^2+^·ADP), and PTMs were apparent from the densities (***Figure 1E, Figure 1-figure supplement 2***). This allowed us to resolve the N-terminus of actin isoforms that is often disordered or missing in prior structures (34). For ß-actin and g-actin, we could resolve the entire N-terminus starting from amino acids D1 and E1, respectively. For skeletal a- and cardiac a-actin, we could resolve the N-terminus starting from amino acids E4 and D2, respectively. The first three amino acids present in mature skeletal muscle actin are unresolved, likely due to their enhanced flexibility in the absence of myosin motors and ABPs (***Figure 1B***). The superposition of all actin isoform structures shows a root-mean-square deviation (RMSD) between 0.83 Å to 1.04 Å, indicating an overall similar topology.

Consistent with the high sequence conservation across actin isoforms (***Figure 1-figure supplement 1B-C***), our reconstructions show the characteristic double-stranded actin helix (***Figure 1A-D, Figure 1-figure supplement 1A***) that has been observed in numerous previous structural studies, including high-resolution cryo-EM studies (16, 19, 25, 29–31, 35, 36). The actin filament itself is composed of G-actin (42 kDa) protomers that are oriented in the same direction (16). Each protomer folds into four subdomains that are referred to as SD1-SD4 (***Figure 1, Figure 1-figure supplement 1A***) (17). SD1 and SD2 form the outer domain, SD3 and SD4 form the inner domain (***Figure 1***). This domain arrangement results in the formation of two clefts – the inner cleft and the outer cleft (2, 14, 37).

The inner cleft between SD2 and SD4 is part of the nucleotide-binding site which is occupied by Mg^2+^·ADP in our structures (***Figure 1E, Figure 2A-D, Figure 1-figure supplement 1A***). The outer cleft between SD1 and SD3 represents a major binding interface for myosins and ABPs (18). It further mediates longitudinal interfaces within the actin filament. SD1 and SD2 of an actin protomer are at the pointed end, SD3 and SD4 are at the barbed end (***Figure 1-figure supplement 1A***) (2, 14, 37). The longitudinal interface between two adjacent actin protomers involves the extended D-loop located in SD2 of one actin protomer that interacts with amino acids located in SD1 and SD3 of another protomer. Both, the N- and C-terminus of the actin protomer are in SD1 (***Figure 1-figure supplement 1A***) (2, 14, 37).

### Similarities and differences between actin isoforms

Actin isoforms differ by conservative and nonconservative substitutions (***Figure 1-figure supplement 1C***) that contribute to their distinct biochemical and *in vivo* function (1, 11, 12). Overall, the amino acid sequence divergence is higher between muscle and nonmuscle actins than within muscle or nonmuscle actin sequences (***Figure 1-figure supplement 1C***). A structural comparison of our cryo-EM reconstructions shows amino acid substitutions across isoforms with the positions of substituted amino acids highlighted (***Figure 3***). Actin isoforms show the largest divergence at the acidic N-terminus within SD1 (***Figure 1E, Figure 1-figure supplement 1C, Figure 3A***) where four conservative substitutions allow for functional interactions with myosins and numerous ABPs (***Figure 1-figure supplement 1C***). Of note, amino acids 1-3 in our structure of skeletal α-actin and the first amino acid in cardiac α-actin are not resolved and therefore not shown in Figure 3. Other substitutions are within SD1 (***Figure 3B, Figure 3-figure supplement 1***), SD3 (***Figure 3C, Figure 3-figure supplement 1***), and SD4 (***Figure 3D, Figure 3-figure supplement 1***). There are no substitutions in SD2, the smallest and most flexible subdomain (34). No significant changes in lateral and longitudinal interfaces and the pitch of the actin helix were observed between actin isoforms, emphasizing their overall conserved filamentous structure in the absence of myosin motors or ABPs (***Figure 3-figure supplement 1***).

**Figure 3:**
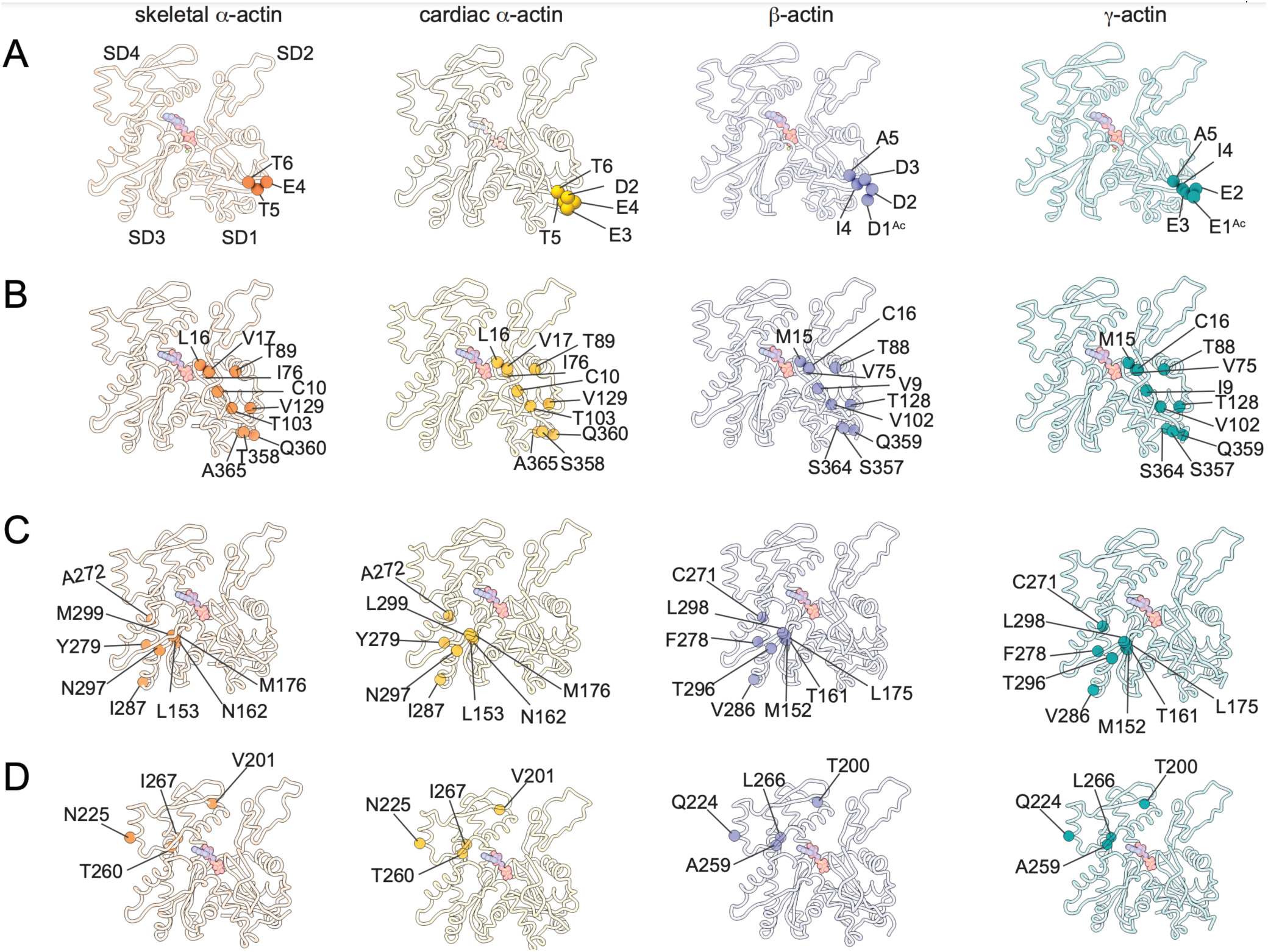
Similarities and differences between actin isoforms. (A) Sequence variations at the N-terminus located in SD1 of actin isoforms. (B) Sequence variations in SD1 of actin isoforms. (C) Sequence variations in SD3 of actin isoforms. (D) Sequence variations in SD4 of actin isoforms. SD2 is conserved between actin isoforms. Substituted amino acids are shown in spheres representation. The identical and nonidentical amino acids at sites of substitutions within the actin protomer across isoforms are shown for skeletal α-actin (orange), cardiac α-actin (yellow), β-actin (purple), and γ-actin (teal) as spheres. Note that the first 3 amino acids of skeletal α-actin and the first amino acid of cardiac α-actin are unresolved in our structures.

Amino acid substitutions at subdomain interfaces, such as the nucleotide-binding site of actin isoforms, are likely to influence protein function. To evaluate their possible impact on nucleotide coordination and the structural organization of the binding site, we performed a comparative structural analysis. Our cryo-EM reconstructions show that the nucleotide-binding site in the inner cleft is highly conserved between actin isoforms (***Figure 1E, Figure 2***). The densities for Mg^2+^ and ADP were assigned without ambiguity and revealed strong interactions with amino acids located in SD2 (Q59, Y69) and SD4 (E207, R210, K213, E214) but also with amino acids located in SD1 (L16/M15, K18, Q137, Y337) and SD3 (D157, M305, Y306, K336) (***Figure 2***). While most of the interactions are polar and electrostatic, amino acid Y306 forms π-π interactions with the adenine ring of ADP. Amino acids L16/M15 and M305/M304 form hydrophobic interactions in the active site (***Figure 2***). The positions of the nucleotide and Mg^2+^ are virtually identical in all structures, suggesting that the substitution of amino acid L16 in muscle actins with M15 in nonmuscle actins does not alter the overall topology of the nucleotide-binding site. Instead, the larger side chain of M16, located in a loop that protrudes into the cleft, acts as an extended lid that flanks the active site and shields the phosphate groups of ADP (***Figure 2***). Near the nucleotide-binding site, amino acids C10 and V17 in muscle actins are substituted with V9 and C16 in nonmuscle actins (***Figure 3, Figure 1-figure supplement 1C***). These substitutions are conservative in that they maintain the overall oxidation-reduction environment within filamentous actin which is important for its dynamic properties and the interaction with some regulatory proteins (38–41).

PTMs are not only essential for actin structure, function, and dynamics but also for their interaction with myosin motors and APBs (42–44). PTMs have been identified on >90 out of the 375 amino acids of actin isoforms (44). Widely documented and critical PTMs of mature actins include the posttranslational acetylation of the N-terminus (∼100%) by N-terminal acetyltransferase NAA80 and methylation of H73 by SET domain protein 3 (SETD3) (43, 45). We could resolve the entire N-terminus including the Nt-acetylated D1 (D1^AC^) in β-actin and the Nt-acetylated E1 (E1^AC^) in γ-actin. To our knowledge, these are the first reported structures of filamentous nonmuscle actins in which the acetylated N-terminus has been resolved. The acetylation site is exposed on the filament surface and adds an additional negative charge to the already negatively charged N-terminus (***Figure 1-figure supplement 1C***). Further, we could resolve the methylated H73 (H73^me^) that contributes to the overall stability of actin isoforms in all cryo-EM reconstructions (46, 47). The presence of both key PTMs in our cryo-EM reconstructions emphasizes that our recombinant human actins represent *bona fide* posttranslationally processed forms of mature nonmuscle actins (***Figure 1B***) (48).

### Myosin modulates actin filament structure

Myosin motors bind actin in a nucleotide-dependent, reversible manner to generate force and motion (49, 50). The myosin enzymatic cycle can be categorized into states with weak (ATP, ADP·Pi) and strong (ADP, nucleotide-free (apo)) actin affinity (51, 52). In the strong affinity states, a binding interface is established between actin and the myosin motor domain (18, 53, 54). To determine whether myosin binding to actin modulates actin filament structure, we compared the structures of filamentous bare actin isoforms with previous high-resolution cryo-EM structures of myosin-bound actins. First, we compared the structure of apo nonmuscle myosin-2C (NM2C, PDB ID: 5JLH) bound to γ-actin, the structure of apo myosin-1B bound to α-skeletal actin (M1B, PDB ID: 6C1H), and the structure of Mg^2+^·ADP.M1B bound to α-skeletal actin (PDB ID: 6C1G) with our structures of bare actin isoforms. The actomyosin structures were selected since they represent distinct high-affinity binding states (Mg^2+^·ADP *versus* apo) and classes of myosin motors with different enzymatic output (24, 52, 55). We found that the binding of myosin to actin causes subtle conformational changes in the actin filament (***Figure 4***). Superimposition of the actin structures revealed that the C-αs of bare and myosin-bound actin protomers deviate with an RMSD of ∼1.15Å. Notably, these changes are similar for myosins from different classes and independent of the nucleotide state of the respective motor domain (***Figure 4***).

**Figure 4:**
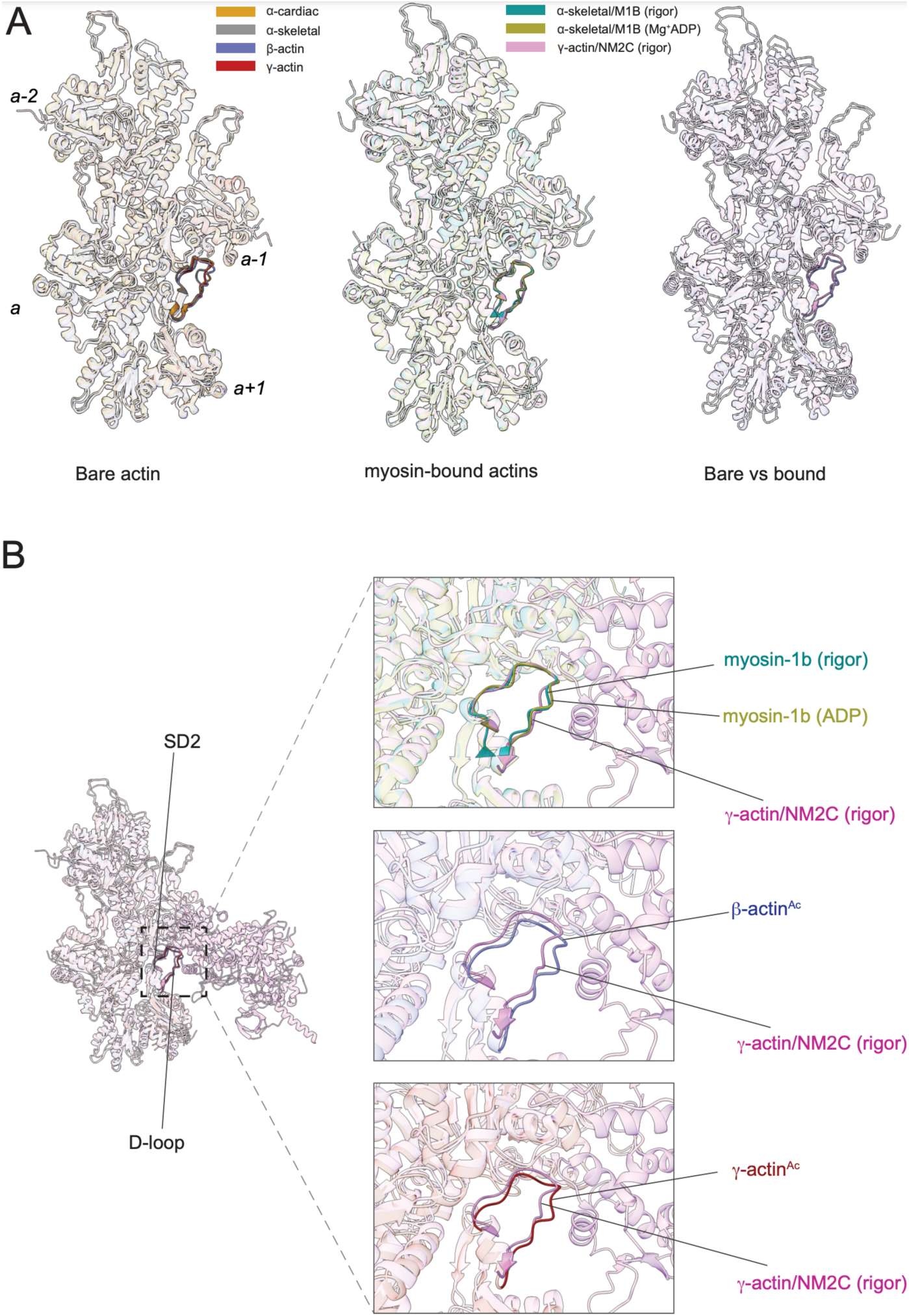
The actomyosin interface. (A) Superimposition of bare actin isoform structures in the Mg^2+^·ADP state (left), superimposition of myosin-bound actin isoforms structures (middle), and overlap of bare vs myosin-bound actin structures (right) are shown. (B) Zoomed-in view of the actomyosin interface at the D-loop region. For clarity, only D-loops involved in the binding of myosins are highlighted in the respective dark colors.

The binding interface between actin and myosin is dynamic. It spans across two adjacent actin protomers along the long-pitch helix of actin and involves SD1 and SD2 of one protomer and the D-loop of another (18, 24). Structural comparison of bare and myosin-bound actins showed that the actin D-loop adopts distinct conformations with an RMSD of ∼0.95-1.25Å (***Figure 4A and*** ***B***). A direct comparison of bare γ-actin with γ-actin/NM2C complex shows the subtle inward movement of the D-loop with an RMSD of ∼1.25Å in the myosin-bound state. This suggests that myosin-binding induces subtle changes in the outer actin cleft that represents a part of the actomyosin interface. Further, the analysis of the actomyosin interface shows that the D-loop conformation does not change significantly with respect to the nucleotide states (apo *versus* ADP) of myosin (***Figure 5B***).

**Figure 5:**
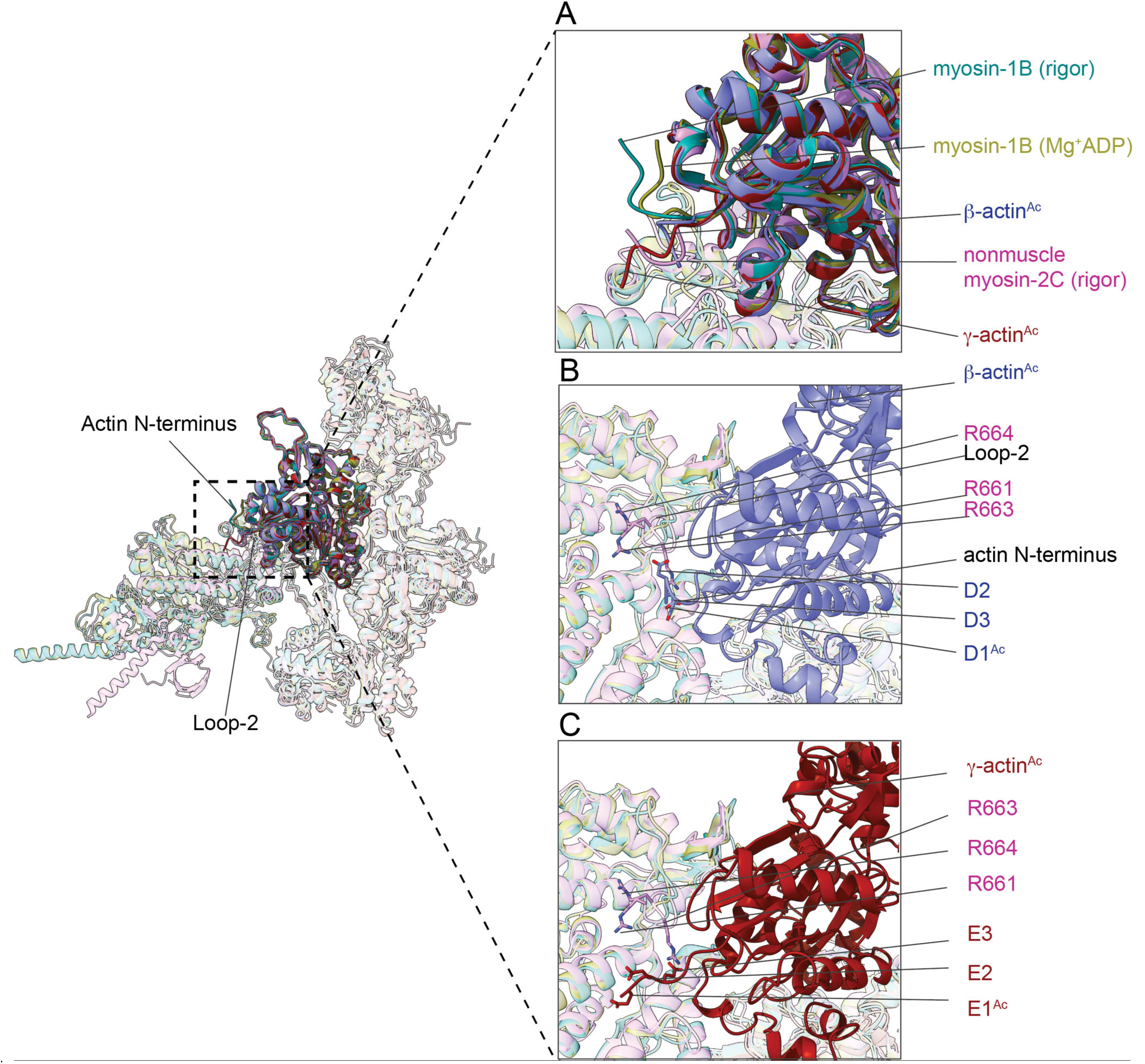
The N-terminus of actin interacts with loop-2 of myosins. (A) Close-up view of the N-termini of bare β-actin (blue) and bare γ-actin (red) and myosin-bound actin isoforms. (B) A cluster of positively charged amino acids in the loop-2 of myosins are in close proximity to amino acids D1^Ac^ to D3 in the N-terminus of bare β-actin (blue) and, (C) a cluster of positively charged amino acids in the loop-2 of myosins are in close proximity to amino acids E1^Ac^ to E3 in the N-terminus of bare γ-actin (red).

Next, we analyzed the position of the exposed actin N-terminus and the effect of Nt-acetylation of nonmuscle actins on the interaction with myosin motors. The structure of NM2C bound to γ-actin (PDB ID: 5JLH) was selected because structural elements involved in actin binding are partially resolved. The superimposition of bare and myosin-bound actin structures shows that the N-termini of skeletal α-actin points in a different direction compared to those from ß-actin and γ-actin (***Figure 5B***). We also observed that the actin N-terminus is positioned closer to the filament surface in myosin-bound structures compared to bare actin structures. The negatively charged N-termini of bare ß-actin and γ-actin are near loop-2, a major element of the actomyosin interface that is rich in positively charged amino acids and variable in length (55–58). This interface is likely to represent one of the initial contact sites upon the formation of a transient intermediate between both proteins during complex formation. Nt-acetylation of actin likely enhances long-range electrostatic interactions between both proteins that subsequently trigger allosteric structural changes that result in the formation of the actomyosin interface. Specifically, a cluster of positively charged amino acids (R661, R663, R664) in loop-2 of NM2C may interact with either the acetylated N-terminus of ß-actin (D1^AC^, D2, D3) or the acetylated N-terminus of γ-actin (E1^AC^, E2, E3) (***Figure 5B-C***) to stabilize the interface. Our structural analysis also revealed that a shorter loop-2, as it is found for example in M1B, might be less efficient in stabilizing the actin N-terminus (***Figure 5***). This suggests that distinct actin-myosin interfaces are formed between actin isoforms and myosin motor proteins that may determine the strength of the interaction and biochemical outputs.

### Amino acid substitutions overlap with the location of mutations and PTMs in actin isoforms

The defining feature of actin isoforms are amino acid substitutions in SD1, SD3, and SD4 (***Figure 3, Figure 1-figure supplement 1C***). Further diversification of actin isoform is achieved by isoform-specific and -redundant PTM mechanisms that contribute to their biochemical and physiological signatures (Table 2). Mutations in genes encoding for actin isoforms have been linked to debilitating diseases including disorders of the muscle and vasculature and cancers (59–62). Abnormal PTM patterns are also often linked to disease and can, among other factors, occur in response to mutations of amino acids that would harbor a PTM under physiological conditions. To identify a possible relationship between the location of mutations and PTMs, we systematically analyzed the location of disease-causing mutations (Table 3) and PTMs (Table 2) in the structures of actin isoforms. First, we mapped select mutations for each actin isoform using our cryo-EM structures. We observed that most mutations are localized in either SD1, SD3, or SD4. Relatively few mutations are localized in SD2 (***Figure 6***), the most conserved subdomain. By comparing the location of mutations between actin isoforms, we found that more mutations are present at the interface along the short-pitch helix, especially in SD3, in muscle actins compared to nonmuscle actins (***Figure 6, Figure 3-figure supplement 1***). PTMs have been reported on ∼25% of the amino acids in actin isoforms with phosphorylation being the most abundant modification (***Table 2***). By comparing the number of reported phosphorylation sites, we found that sites in nonmuscle actins are more often modified than sites in muscle actins. In some instances, phosphorylation is abolished by amino acid substitutions between actin isoforms. For instance, the phosphorylatable T161 and T200 in nonmuscle actins are substituted with non-phosphorylatable amino acids in muscle actins (***Table 2***). Similarly, the substitution of C271 in nonmuscle actins with A272 in muscle actins changes the oxidation-reduction environment between actin isoforms (***Table 2***). This suggests that an isoform-specific ‘PTM code’ exists that is likely to change in time and space as cellular conditions demand. By mapping the precise location of PTMs, we found that isoforms undergo extensive post-translational modifications throughout the molecule except for the regions corresponding to amino acids 125–145, 240–256, and 331–354, which have an overall lower PTM burden (***Table 2).*** This region includes the pathogenic helix (K112-T126) that mediates the interaction between two actin protomers (44). Only two amino acids (K112/K113 and K117/K118) have been described to be posttranslationally modified while this region is known as a mutational hotspot in actin isoforms (***Figure 6 and Table 3***). As structural changes in the pathogenic helix and structural elements that interact with it can affect filament biochemistry, dynamics, and interactions with binding proteins, altered PTM patterns and/or mutations in this region could underlie one of the molecular mechanisms that lead to disease (44, 63, 64).

**Figure 6:**
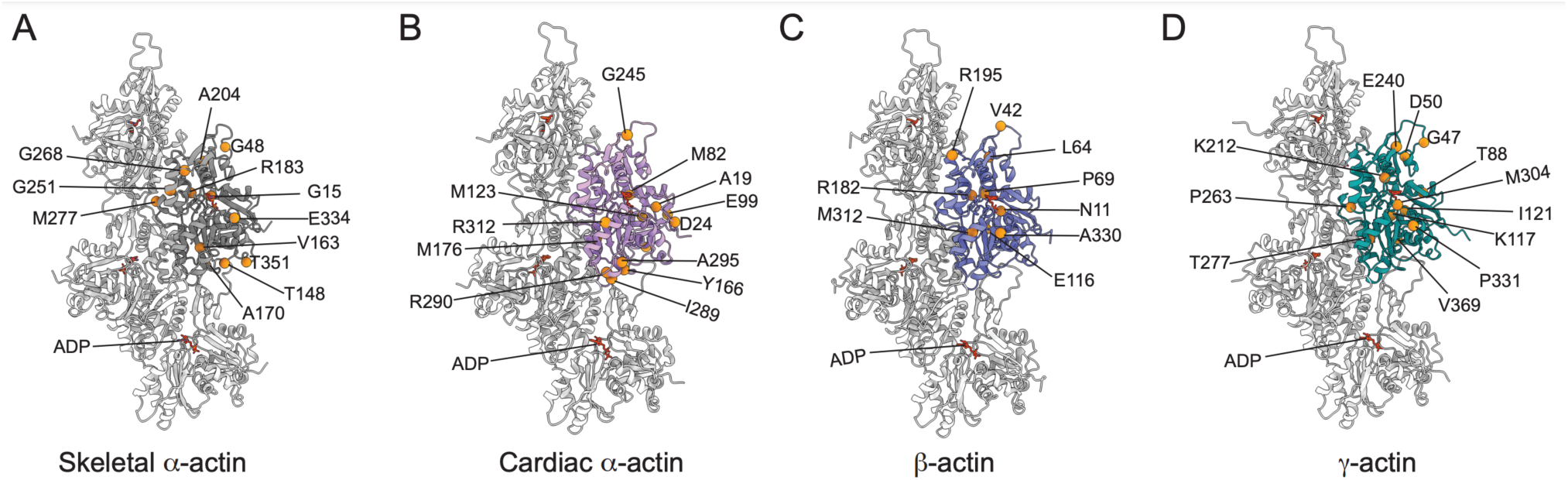
Location of selected mutations in actin isoform structures. (A) Select mutations (n > 4) in skeletal α-actin are mapped onto the α-actin structure. (B) Select mutations (n > 4) in cardiac α-actin are mapped onto the cardiac α-actin structure. (C) Select mutations (n > 2) in cytoplasmic β-actin are mapped onto the acetylated β-actin structure and (D) Select mutations (n > 4) in cytoplasmic γ-actin are mapped onto the acetylated γ-actin structure. ‘n’ indicates the number of studies that showed mutations of the indicated amino acids.

**Table 2:** Posttranslational modifications of mammalian actin isoforms (41, 44, 81). ^a^ denotes amino acid sites with multiple PTMs.

| **Nonmuscle actins ($\beta/\gamma$-actin)** |  |
| --- | --- |
| **Actin PTM** | **Modified amino acids** |
| Acetylation | D1, E1, K49<sup>a</sup>, K60<sup>a</sup>, K67<sup>a</sup>, K112<sup>a</sup>, K190<sup>a</sup>, K212<sup>a</sup>, K314<sup>a</sup>, K325<sup>a</sup>, K327<sup>a</sup> |
| ADP-ribosylation | T147<sup>a</sup>, R176 |
| Arginylation | D2, S51<sup>a</sup>, F89 |
| Carbonylation | H39, H86, H172, C373<sup>a</sup> |
| Crosslinking | K49/E269<sup>a</sup> |
| Disulfide bond | C284<sup>a</sup>, C373<sup>a</sup> |
| Glutathionylation | C216<sup>a</sup>, C373<sup>a</sup> |
| Isoaspartylation | D24, D178 |
| Methylation | K17<sup>a</sup>, K67<sup>a</sup>, H72, K83<sup>a</sup> |
| Tyrosine nitration | Y52<sup>a</sup>, Y68<sup>a</sup>, Y90<sup>a</sup>, Y197<sup>a</sup>, Y217<sup>a</sup>, Y239<sup>a</sup>, Y293<sup>a</sup>, Y361<sup>a</sup> |
| S-nitrosylation | C216<sup>a</sup>, C256<sup>a</sup>, C284<sup>a</sup>, C373<sup>a</sup> |
| Oxidation | C16, M43, M46, M81, M189, C216<sup>a</sup>, M226, C256<sup>a</sup>, M268, C271, C284<sup>a</sup>, M324, M354, C373<sup>a</sup> |
| Phosphorylation | S13, S32, S51<sup>a</sup>, Y52<sup>a</sup>, S59, T65, Y68<sup>a</sup>, T76, T88, Y90<sup>a</sup>, Y142, T147<sup>a</sup>, S154, T159, T161, Y165, Y168, T185, Y197<sup>a</sup>, S198, T200, T201, T202, Y217<sup>a</sup>, T228, S232, S234, S238, Y239<sup>a</sup>, T248, S264, S270, Y293<sup>a</sup>, T296, S299, Y305, T317, S322, T323, Y361<sup>a</sup>, S364 |
| SUMOylation | K60<sup>a</sup>, K67<sup>a</sup>, K83<sup>a</sup>, K112<sup>a</sup>, K283<sup>a</sup>, K290<sup>a</sup>, K314<sup>a</sup>, K325<sup>a</sup>, K327<sup>a</sup> |
| Transglutamination | Q40 |
| Ubiquitination | K17<sup>a</sup>, K49<sup>a</sup>, K60<sup>a</sup>, K67<sup>a</sup>, K83<sup>a</sup>, K112<sup>a</sup>, K190<sup>a</sup>, K212<sup>a</sup>, K214, K237, K283<sup>a</sup>, K290<sup>a</sup>, K314<sup>a</sup>, K325<sup>a</sup>, K327<sup>a</sup>, K358 |
| **Muscle actins (cardiac α-actin, smooth α-actin, skeletal α-actin, and smooth γ-actin). Residues are numbered based on the sequence of skeletal α-actin.** |  |
| **Actin PTM** | **Modified amino acid** |
| Acetylation | D1, K50<sup>a</sup>, K61<sup>a</sup>, K84<sup>a</sup>, K113<sup>a</sup>, K191<sup>a</sup>, K213, K215<sup>a</sup>, K315<sup>a</sup>, K326<sup>a</sup>, K328<sup>a</sup> |
| Arginylation | S52<sup>a</sup>, I85<sup>a</sup>, G150, L293, N297<sup>a</sup> |
| Malonylation | K61<sup>a</sup> |
| Methylation | K68<sup>a</sup>, K84<sup>a</sup>, I85<sup>a</sup>, R95, K215<sup>a</sup>, R254, N297<sup>a</sup> |
| Oxidation | W79, W86, M176, W340, W356 |
| Phosphorylation | S52<sup>a</sup>, S60, T66, Y69, T77, T89, Y143, Y166, Y169, T186, Y188, T194, Y198, S199<sup>a</sup>, Y218, S233, S234, S239, Y240, T249, T260, S265, Y279, Y294, T304, Y306, T318, S323<sup>a</sup>, T324, Y362 |
| O-GlcNAcylation | S52<sup>a</sup>, S155, S199<sup>a</sup>, S232, S323<sup>a</sup>, S368 |
| SUMOylation | K61<sup>a</sup>, K315<sup>a</sup>, K326<sup>a</sup>, K328<sup>a</sup> |
| Transglutamination | Q41 |
| Ubiquitination | K50<sup>a</sup>, K61<sup>a</sup>, K68<sup>a</sup>, K84<sup>a</sup>, K113<sup>a</sup>, K191<sup>a</sup>, K238, K291, K315<sup>a</sup>, K326<sup>a</sup>, K328<sup>a</sup> |

**Table 3:** Location of selected disease-causing mutations in actin isoforms (60, 82–84). *Denotes termination.

| Skeletal α- actin | | Cardiac α- actin | | β-actin | | γ-actin | |
| --- | --- | --- | --- | --- | --- | --- | --- |
| Amino acid | Mutation | Amino acid | Mutation | Amino acid | Mutation | Amino acid | Mutation |
| 1 | D1Y | 1 | D1H | 11 | N11D,H | 11 | N11D |
| 4 | E4K | 8 | L8M | 42 | V42M | 31 | P31S |
| 11 | D11N | 19 | A19V | 46 | M46T | 39 | H39Y |
| 15 | G15R,S,D | 21 | F21L | 58 | Q58R | 47 | G47R |
| 25 | D25N | 24 | D24N | 64 | L64F,V | 50 | D50N |
| 35 | V35L,A | 26 | A26V | 69 | P69A,L | 57 | A57V |
| 36 | G36A | 35 | V35A | 72 | H72D | 65 | T65I |
| 38 | P38L | 40 | H40Y | 73 | G73S | 69 | P69L |
| 39 | R39H,* | 46 | G46S | 74 | I74T | 74 | I74L |
| 40 | H40Y | 47 | M47L | 102 | V102L | 81 | M81L |
| 41 | Q41R | 63 | G63D | 116 | E116D,K | 88 | T88I |
| 42 | G42V | 75 | I75V | 118 | M118T | 117 | K117N,M |
| 43 | V43F | 76 | I76N | 119 | T119I | 119 | T119I |
| 44 | M44T | 82 | M82T | 148 | T148I | 121 | I121V |
| 46 | G46D,C | 88 | H88Y | 161 | T161A | 134 | A134V |
| 47 | M47V | 91 | Y91C,H | 170 | L170F | 144 | S144C |
| 48 | G48A,C,D | 92 | N92S | 178 | D178E | 152 | M152I |
| 55 | G55R | 95 | R95C | 182 | R182Y | 154 | S154F |
| 64 | I64N,S | 99 | E99K | 195 | R195C,H | 166 | E166K |
| 66 | T66N,I | 101 | H101Q | 203 | A203G | 178 | D178Y |
| 68 | K68R | 108 | A108V | 205 | R205Q | 180 | A180G |
| 70 | P70R | 119 | M119V | 208 | V208L | 182 | R182Q |
| 71 | I71F,V | 123 | M123V | 243 | G244S | 186 | D186H |
| 72 | E72K | 126 | T126I | 267 | G267R | 191 | I191F |
| 73 | H73R,N,L | 157 | D157N | 312 | M312R | 202 | T202K,M |
| 74 | G74D | 164 | P164A | 330 | A330V | 209 | R209C |
| 75 | I75L,S | 166 | Y166C | 363 | E363K | 212 | K212R |
| 77 | T77A | 170 | A170T | 364 | S367L | 240 | E240K |
| 83 | E83K | 173 | H173R | 372 | K372* | 242 | P242L |
| 92 | N92K | 176 | M176L |  |  | 253 | R253W |
| 94 | L94P | 206 | R206H |  |  | 255 | R255W |
| 107 | E107D | 210 | R210H |  |  | 257 | P257L |
| 111 | N111S | 218 | Y218H,S |  |  | 263 | P263L |
| 112 | P112L | 222 | D222Y |  |  | 267 | G267S |
| 113 | K113E | 229 | T229R |  |  | 277 | T277I |
| 114 | A114T,V | 230 | A230T,V |  |  | 282 | M282T,V |
| 115 | N115T,S | 234 | S234F |  |  | 304 | M304T |
| 116 | R116H | 245 | G245D |  |  | 315 | E315K |
| 117 | E117Q | 249 | T249S |  |  | 324 | M324K |
| 120 | T120S | 250 | I250M,T |  |  | 331 | P331A,S |
| 132 | M132V | 263 | Q263E |  |  | 333 | E333Q |
| 134 | V134A | 264 | P264L |  |  | 334 | R334H |
| 136 | I136M,T | 267 | I267T |  |  | 348 | L348M |
| 137 | Q137H | 271 | S271F |  |  | 369 | V369A |
| 138 | A138D,P | 282 | I282F |  |  |  |  |
| 139 | V139A | 289 | I289T |  |  |  |  |
| 140 | L140P | 290 | R290G |  |  |  |  |
| 142 | L142F | 294 | Y294H,N |  |  |  |  |
| 143 | Y143* | 295 | A295S |  |  |  |  |
| 144 | A144V | 305 | M305L |  |  |  |  |
| 146 | G146D,S | 311 | D311H |  |  |  |  |
| 147 | R147K | 312 | R312C,H |  |  |  |  |
| 148 | T148A,I,N,S | 321 | A321V |  |  |  |  |
| 150 | G150A | 327 | I327T |  |  |  |  |
| 152 | V152L | 329 | I329N |  |  |  |  |
| 154 | D154N | 355 | M355V |  |  |  |  |
| 158 | G158C | 361 | E361G |  |  |  |  |
| 161 | H161D | 365 | A365T |  |  |  |  |
| 163 | V163L,M | 369 | I369T |  |  |  |  |
| 170 | A170E,G |  |  |  |  |  |  |
| 178 | L178P |  |  |  |  |  |  |
| 179 | D179N,G,H |  |  |  |  |  |  |
| 181 | A181T |  |  |  |  |  |  |
| 182 | G182D |  |  |  |  |  |  |
| 183 | R183C,G,S |  |  |  |  |  |  |
| 184 | D184G,H |  |  |  |  |  |  |
| 188 | Y188* |  |  |  |  |  |  |
| 189 | L189P |  |  |  |  |  |  |
| 191 | K191N |  |  |  |  |  |  |
| 194 | T194P |  |  |  |  |  |  |
| 195 | E195D |  |  |  |  |  |  |
| 196 | R196H,L |  |  |  |  |  |  |
| 197 | G197S |  |  |  |  |  |  |
| 198 | Y198C |  |  |  |  |  |  |
| 202 | T202I |  |  |  |  |  |  |
| 204 | A204T |  |  |  |  |  |  |
| 205 | E205D,G |  |  |  |  |  |  |
| 215 | K215* |  |  |  |  |  |  |
| 221 | L221P |  |  |  |  |  |  |
| 224 | E224Q,G |  |  |  |  |  |  |
| 226 | E226Q |  |  |  |  |  |  |
| 227 | M227I,T,V |  |  |  |  |  |  |
| 230 | A230V |  |  |  |  |  |  |
| 237 | E237* |  |  |  |  |  |  |
| 241 | E241K |  |  |  |  |  |  |
| 244 | D244E |  |  |  |  |  |  |
| 245 | G245R |  |  |  |  |  |  |
| 246 | Q246R,K |  |  |  |  |  |  |
| 250 | I250T |  |  |  |  |  |  |
| 251 | G251D,R |  |  |  |  |  |  |
| 252 | N252Y |  |  |  |  |  |  |
| 253 | E253G |  |  |  |  |  |  |
| 254 | R254H |  |  |  |  |  |  |
| 255 | F255C |  |  |  |  |  |  |
| 256 | R256C,H,L |  |  |  |  |  |  |
| 259 | E259V |  |  |  |  |  |  |
| 263 | Q263L |  |  |  |  |  |  |
| 264 | P264T |  |  |  |  |  |  |
| 265 | S265C |  |  |  |  |  |  |
| 266 | F266L |  |  |  |  |  |  |
| 268 | G268R,D,C |  |  |  |  |  |  |
| 269 | M269R,V |  |  |  |  |  |  |
| 270 | E270Q |  |  |  |  |  |  |
| 272 | A272E,V |  |  |  |  |  |  |
| 279 | Y279H |  |  |  |  |  |  |
| 280 | N280K |  |  |  |  |  |  |
| 283 | M283K,R |  |  |  |  |  |  |
| 286 | D286G |  |  |  |  |  |  |
| 288 | D288N,H |  |  |  |  |  |  |
| 289 | I289F |  |  |  |  |  |  |
| 292 | D292V |  |  |  |  |  |  |
| 295 | A295T |  |  |  |  |  |  |
| 299 | M299K |  |  |  |  |  |  |
| 306 | Y306C |  |  |  |  |  |  |
| 323 | S323R |  |  |  |  |  |  |
| 325 | M325K |  |  |  |  |  |  |
| 326 | K326N |  |  |  |  |  |  |
| 332 | P332R,S |  |  |  |  |  |  |
| 334 | E334A,K |  |  |  |  |  |  |
| 336 | K336E,I,T |  |  |  |  |  |  |
| 346 | L346Q |  |  |  |  |  |  |
| 348 | S348L |  |  |  |  |  |  |
| 351 | T351A |  |  |  |  |  |  |
| 352 | F352S |  |  |  |  |  |  |
| 356 | W356C |  |  |  |  |  |  |
| 357 | I357L |  |  |  |  |  |  |
| 367 | P367L |  |  |  |  |  |  |
| 369 | I369L |  |  |  |  |  |  |
| 370 | V370F |  |  |  |  |  |  |
| 372 | R372C,S |  |  |  |  |  |  |
| 373 | K373E,Q,N |  |  |  |  |  |  |
| 374 | C374S |  |  |  |  |  |  |
| 375 | F375C,Y |  |  |  |  |  |  |

## Discussion

Despite many elegant, high-resolution cryo-EM studies of filamentous actin in the absence and presence of myosin motors and ABPs, our understanding of the structural mechanisms that underlie the nuanced interactions of actin isoforms with interacting proteins remain largely elusive. This knowledge however is critical to understanding how protein binding modulates actin filament structure that in turn may be recognized by and guide other interacting proteins or partners (9, 10, 25). That way, specialized actin networks with distinct structures and compositions can form and a variety of actin landscapes can be created (9, 10, 65–68).

To further our understanding of the sequence-structure relationship of actin isoforms, we solved four high-resolution cryo-EM structures including the structure of bare and homogeneous filamentous skeletal α-actin and the previously unknown structures of cardiac α-actin, ß-actin, and γ-actin. This allowed us to reveal the similarities and differences between actin isoforms, their distinct modes of interactions with myosin motors, as well as the analysis of the complex interplay between disease-causing mutations and the ‘PTM code’. Our study showed that the overall structure of actin isoforms is conserved, as expected from the high sequence conservation. However, each of these structures shows subtle differences. Minor changes were observed near the nucleotide-binding cleft (***Figure 2***) that may contribute to the reported changes in the biochemistries of actin isoforms (69, 70). The most prominent structural change between actin isoforms corresponds to the N-terminus location (***Figure 1, Figure 5***). Given that the sequence of actin isoforms differs the most at the N-terminus (***Figure 1E, Figure 1-figure supplement 1C***) and the central role of the N-terminus in interactions with myosin motors and ABPs, we propose that these topological changes may drive different biochemical interactions. For example, by comparing the structures of α-skeletal actin with the γ-actin/NM2C complex, a previous study suggested a pulling mechanism in which the actin N-terminus is pulled towards the actomyosin interface in the rigor state (18). Based on our structures that show different conformations of the N-terminus, we suggest that instead of a large conformational change through a pulling mechanism, actin isoform-specific interfaces with myosin motors are formed that involve a more subtle movement of the N-terminus due to retropropagated structural changes. These interfaces likely contribute to the extent actin isoforms can activate the myosin ATPase activity and support motility *in vitro* and establish different actin landscapes in cells (7, 10). Importantly, these actin isoform-specific interfaces may be further diversified by differential N-terminal processing of actin isoforms, PTMs, and alternatively spliced myosin motors (7, 15, 41, 44, 55, 71, 72). For example, the Nt-acetylated N-termini of actin isoforms would likely be more efficient to establish electrostatic interactions with the positively charged loop-2 of myosin while the addition of a positive charge to the N-terminus of actin isoforms through arginylation may weaken the interaction. While we focused our structural analysis on the interaction between actin isoforms with myosin motors, the actin N-terminus can interact with numerous ABPs (15). Therefore, we predict that the proposed isoform-specific interaction mechanisms extend to ABPs. Together, we present direct structural evidence that the binding of myosin motors modulates actin filament structure and propose a possible mechanism for the formation of actin isoform-specific interactions with binding proteins by the conformation of its N-terminus (***Figure 5***).

Actin isoforms are subject to extensive PTM mechanisms that may drive structural changes, distinct biochemistries, and cellular activities (3, 12, 41). Prevalent PTMs of actin isoforms include acetylation, phosphorylation, and methylation (Table 2). The high resolution of our structures allowed us to resolve the Nt-acetylated N-termini in filamentous ß-actin and γ-actin, to our knowledge, for the first time. Further, we could resolve the methylated H73/H72 in all our structures. By mapping the known locations of PTMs (***Table 2***) and disease-causing mutations (***Table 3***) on our structures of actin isoforms, we revealed that some sites of modification overlap with sites of mutation (***Table 2, Table 3, Figure 6***). This suggests that some mutations may not only change the structure and biochemistry of actin isoforms but may elicit complex combinatorial effects by abolishing native posttranslational modification patterns or by potentially promoting aberrant posttranslational modifications. For instance, T88 in β- and γ-actins are part of the binding interfaces with cofilin and gelsolin 3 and can be phosphorylated under normal conditions (44). Mutation of amino acid T88 to I88 in γ-actin not only prevents phosphorylation but has also been linked to age-related hearing loss (73). Another example is amino acid K212 that interacts with nucleotides in the active site (***Figure 2***) and is known to be acetylated and ubiquitinated in β- and γ-actins while mutation K212R has been linked to hearing loss (74). Further, our analysis revealed that in addition to the absence of amino acid substitutions in SD2, its mutation burden is also reduced (***Figure 1-figure supplement 2***). However, PTMs are present in SD2 of actin isoforms, suggesting that major structural alterations induced by amino acid substitutions and/or mutations in the highly conserved D-loop are likely to interfere with its critical role in filament biochemistry and the interactions with ABPs and myosin motors. In summary, our work suggests that in some instances, mutational changes may alter or re-write the ‘PTM code’ of actin isoforms that may further modulate the possible effect of the mutation on protein function.

In conclusion, we present direct evidence for the structural divergence of actin isoforms that underlies their nuanced interactions with myosin motors and ABPs. Our work serves as a strong foundation for our understanding of the sequence-function relationship of actin isoforms. By adding our cryo-EM structures to the collection of previous structures of the bare and decorated actin filaments, we provide a comprehensive understanding of the remarkable diversity of actin isoforms that is reflected in their biological activities in health and disease.

## Material and Methods

### Protein production and purification

#### Native actins

Rabbit skeletal muscle α-actin (UniProt ID: P68135) was prepared from acetone powder (Pel-Freez Biologicals, Rogers, AR) as described earlier (75). Cardiac α-actin (UniProt ID: Q3ZC07) was prepared from the left ventricle of a bovine heart (local butcher) as described for α-skeletal muscle actin. At the amino acid level, both proteins are identical to the respective human proteins.

#### Recombinant actins

*P. pastoris* transformants for human β-actin (UniProt ID: P60709) and human γ-actin (UniProt ID: P63261) were stored at -80°C and were revived on YPD solid media plates at 30°C. Cells were inoculated into 200 ml MGY liquid media composed of 1.34% yeast nitrogen base without amino acids (Millipore Sigma, St. Louis, MO), 0.4 mg/L biotin, and 1% glycerol and cultured at 30°C, 220 rpm. The culture medium was diluted to 6 L with fresh MGY media and cells were further cultured at 30°C, 220 rpm in six 2 L flasks until the optical density at 600 nm (OD600) reached around 1.5. Cells were pelleted down by centrifugation (10628 g at 25°C for 5 min, F9-6 x 1000 LEX rotor, Thermo Fisher Scientific, Waltham, MA). The cells were washed once with sterilized water and re-suspended into 6 L MM composed of 1.34% yeast nitrogen base without amino acids (SIGMA Y0626), 0.4 mg/L biotin and 0.5% methanol. Cells were cultured in twelve 2 L baffled flasks (500 mL for each) at 30°C, 220 rpm for 1.5-2 days. 0.5 % methanol was fed every 24 hours during the culture. Cells were pelleted down by centrifugation (10628 g at 25°C for 5 min, F9-6 x 1000 LEX rotor, Thermo Fisher Scientific, Waltham, MA). Cells were washed once with water and suspended in 75 mL of ice-cold water. The suspension was dripped into a liquid nitrogen bath and stored at -80°C. 50 g cell suspension was loaded into a grinder tube SPEX® SamplePrep, Metuchen, NJ) pre-cooled with liquid nitrogen. Cells were grinded in a liquid nitrogen bath of a Freezer mill (SPEX® SamplePrep, Metuchen, NJ). The duration of the grinding was 1 min with 14 cycle per second (cps). The grinding procedure was repeated 30 times at 1 min intervals. Liquid nitrogen was re-filled every 10 times of the grinding. The resulting powder was kept on dry ice until the next step. The powder was kept at room temperature until it started to melt. Then it was resolved in equal amount of 2 x binding buffer composed of 20mM imidazole (pH 7.4), 20mM HEPES (pH 7.4), 0.6M NaCl, 4mM MgCl_2_, 2mM ATP (pH 7.0), 2x concentration of protease inhibitor cocktail (Roche cOmplete, EDTA free, Millipore Sigma, St. Louis, MO), 1mM phenylmethylsulfonyl fluoride (PMSF) and 7mM beta-mercaptoethanol. The lysate was sonicated on ice (3 minutes, 5 seconds pulse, 10 seconds pause with 60% amplitude, QSONICA SONICATORS, Newtown, CT) until all aggregates were resolved. The lysate was centrifuged at 4°C (3220 g for 5 min, A-4-81 rotor, Eppendorf, Enfield, CT) to remove intact cells and debris. The insoluble fraction was removed by high-speed centrifugation at 4°C (25658 g for 30 minutes, A23-6 x 100 rotor, (Thermo Fisher Scientific, Waltham, MA). The supernatant was filtrated with a Filtropur BT50 0.2 µm bottle top filter (SARSTEDT, Nümbrecht, Germany) and incubated with 6 ml Nickel resin (Thermo Fisher Scientific, Waltham, MA) at 4°C for 1 hour. The resin was pelleted down by centrifugation at 4°C (1258 g for 5 min, A-4-81 rotor, Eppendorf, Enfield, CT) and washed with ice-cold 50 mL Binding buffer composed of 10mM imidazole (pH 7.4), 10mM HEPES (pH 7.4), 300mM NaCl, 2mM MgCl_2_, 1mM ATP (pH 7.0) and 7mM beta-mercaptoethanol for 4 times. The resin was washed 5 times with G-buffer composed of 5mM HEPES (pH 7.4), 0.2mM CaCl_2_, 0.01 w/v% NaN_3_, 0.2mM ATP (pH 7.0) and 0.5mM dithiothreitol (DTT). The resin was suspended in an ice-cold 40 mL G-buffer with 5 µg/ml TLCK treated chymotrypsin (Millipore Sigma, St. Louis, MO) and incubated overnight at 4°C. The chymotrypsin was inactivated by 1 mM PMSF and the elution was collected into a tube. Actin retained on the resin was eluted with 12 mL G-buffer without actin and all elution fractions were combined and concentrated with a 30 kDa cut-off membrane (Millipore Sigma, St. Louis, MO) to 0.9 ml. The 0.9 ml of each actin was polymerized by adding 100 µl of 10x MKE solution composed of 20mM MgCl_2_, 50mM ethylene glycol tetraacetic acid (EGTA), and 1M KCl for 1 hour at room temperature. The polymerized actin samples were pelleted down by ultracentrifugation at room temperature (45000 rpm for 1 hour, TLA-55 rotor, Beckman Coulter, Indianapolis, IN). The pellets were rinsed once with a 1 mL G-buffer and re-suspended into ice-cold 0.5 mL G-buffer. The actin was depolymerized by dialysis against a 1 L G-buffer at 4°C for 2 days. The dialysis buffer was exchanged every 12 hours. The solutions with the depolymerized actin were collected into 1.5 ml centrifuge tubes and stored on ice.

### Sample preparation and Cryo-EM data collection

For cryo-EM studies, G-actin was polymerized by the addition of a 10x stock solution of buffer containing 0.5M KCl, 20mM MgCl_2_, 10mM EDTA, and 0.1M MOPS pH 7.0 before plunge freezing. Briefly, C-flat 1.2/1.3 gold grids were glow-discharged with a PELCO easiGlow (TedPella, Redding, CA) for 60s. 4 µL of actin sample was applied to the grids, incubated for 1 minute, and blotted for 4s at 95% humidity. Grids were plunged into liquid ethane using a Leica EM GM2 plunger (Leica Microsystems, Wetzlar, Germany) and stored in liquid nitrogen until data collection. All grids were screened for homogeneous sample distribution and optimal ice thickness on a Glacios cryo-TEM (Thermo Fisher Scientific, Waltham, MA) equipped with a Falcon 3EC direct electron detector at a magnification of 92,000x. Data collection on optimal grids was performed on a Titan Krios G3i (Thermo Fisher Scientific, Waltham, MA) operated at 300kV, equipped with a K3 direct electron detector, a Bioquantum energy filter, and a Cs image corrector. For skeletal α-actin, a total of 2,046 movies with a magnification of 81,000x, corresponding to a pixel size of 0.4455 Å were collected in super-resolution mode at a defocus range of -0.5 µm to -2.5 µm with a total electron dose of 65 e^-^/Å^2^ per movie. For cardiac α-actin, a total of 1,444 movies with a magnification of 81,000x, corresponding to a pixel size of 0.4455 Å were collected in super-resolution mode at a defocus range of -0.5 µm to -2.5 µm with a total electron dose of 60 e^-^/Å^2^ per movie. For acetylated β- and γ-actins, a total of 1,352 movies and 2,952 movies with a magnification of 81,000x, corresponding to a pixel size of 0.4455 Å were collected in super-resolution mode at a defocus range of -0.5 µm to -2.5 µm with a total electron dose of 50 e^-^/Å^2^ and 65 e^-^/Å^2^ per movie, respectively. Data collection for all actins were performed using EPU software (Thermo Fisher Scientific, Waltham, MA). Data collection statistics are shown in Table 1.

### Image processing and 3D reconstruction

All raw movies of skeletal and cardiac α-actin were aligned, drift corrected, and dose weighted using the Patch motion module and the CTF parameters were estimated using the Patch CTF module implemented in cryoSPARCv3.2 (76). All raw movies of acetylated β- and γ-actin were aligned, and motion-corrected using the Patch motion module in cryoSPARC and MotionCor2 software (76), and the CTF parameters were estimated using the Patch CTF module implemented in cryoSPARCv3.2. Segments of actin filaments were initially picked using the template-free tracing method implemented in the filament tracer module in cryoSPARC to generate templates for particle picking (76). A small set of particles for reference-free 2D classifications was selected and subsequently used as templates for filament tracing. Further, segments of filaments were extracted and several rounds of 2D classifications were performed to remove erroneous picks and well-aligned 2D classes were selected as templates for ab initio 3D reconstruction. For all actins, a major class with a filamentous model that showed clear helical features was selected for 3D refinements using the helical refinement module implemented in cryoSPARCv3.2 (76). For acetylated β- and γ-actins, final rounds of refinements were performed using the homogenous refinement module in cryoSPARC. Data collection statistics, the image processing and the refinement summary of models are shown in Table 1.

### Model building, refinement, and validation

For model building, sharpened maps were used to build models of filamentous actins. We have used a common approach of model building, refinement, and validation for all actin isoforms. Briefly, the high-resolution structures of Mg^2+^.ADP bound skeletal and cardiac α-actins, Nt-acetylated β- and γ-actins were built using PDB entry 6DJO as a template. Initial rigid-body docking into the cryo-EM reconstructions of actin maps was performed with Molecular Dynamics Flexible Fitting (MDFF) function in Chimera (77). Iterative model building was performed using real-space refinement in Phenix (78) and COOT (79). The acetylated N-terminus of the β- and γ-actins were manually placed in the cryo-EM densities using COOT. It is important to note that the N-terminal amino acids (1 to 3) showed high flexibility and we manually placed the best possible rotamers using COOT and minimized the rotamers using Phenix. All final actin models were refined using Phenix and the refined models were validated using MolProbity (80). Amino acids are numbered according to the sequence of mature actin isoforms (***Figure 1-figure supplement 1C***).

## Acknowledgments

We thank Dr. Jim Sellers (National Heart, Lung, and Blood Institute, National Institutes of Health) and Dr. Earl Homsher (University of California, Los Angeles) for bovine α-cardiac actin powder. Electron microscopy was performed at the Center for Electron Microscopy and Analysis (CEMAS) at The Ohio State University. We thank the Ohio Supercomputer Center (OSC) for high-performance computing resources. This work was supported by the National Institute of General Medical Sciences of the National Institutes of Health under Award Number R01GM143539 (KC), the National Heart Lung and Blood Institute of the National Institutes of Health under Award Number K22HL131869 (SMH), and Wellcome Trust (203276/Z/16/Z), European Research Council (ERC-2014-ADG No. 671083) and Biotechnology and Biosciences Research Council (BB/S003789/1) to MKB.

## Author contributions

KC conceived the project and experiments. ASA, HLH, RS, AS, TH, and SMH expressed and/or purified proteins. ASA, YN, and KC carried out cryo-EM experiments. ASA, SMH, and KC analyzed the cryo-EM data. SMH and KC wrote the manuscript with inputs from ASA, HLH, YN, and MKB.

## Data availability

All the structures and electron density maps generated have been deposited in the Protein Data Bank (PDB) and Electron Microscopy Data Bank (EMDB). The PDB and EMDB entries for skeletal α-actin are 8DMX and EMD-27548; for cardiac α-actin 8DMY and EMD-27549; for β-actin 8DNH and EMD-27572; for γ-actin 8DNF and EMD-27565.

---

**Figure 1-figure supplement 1:**
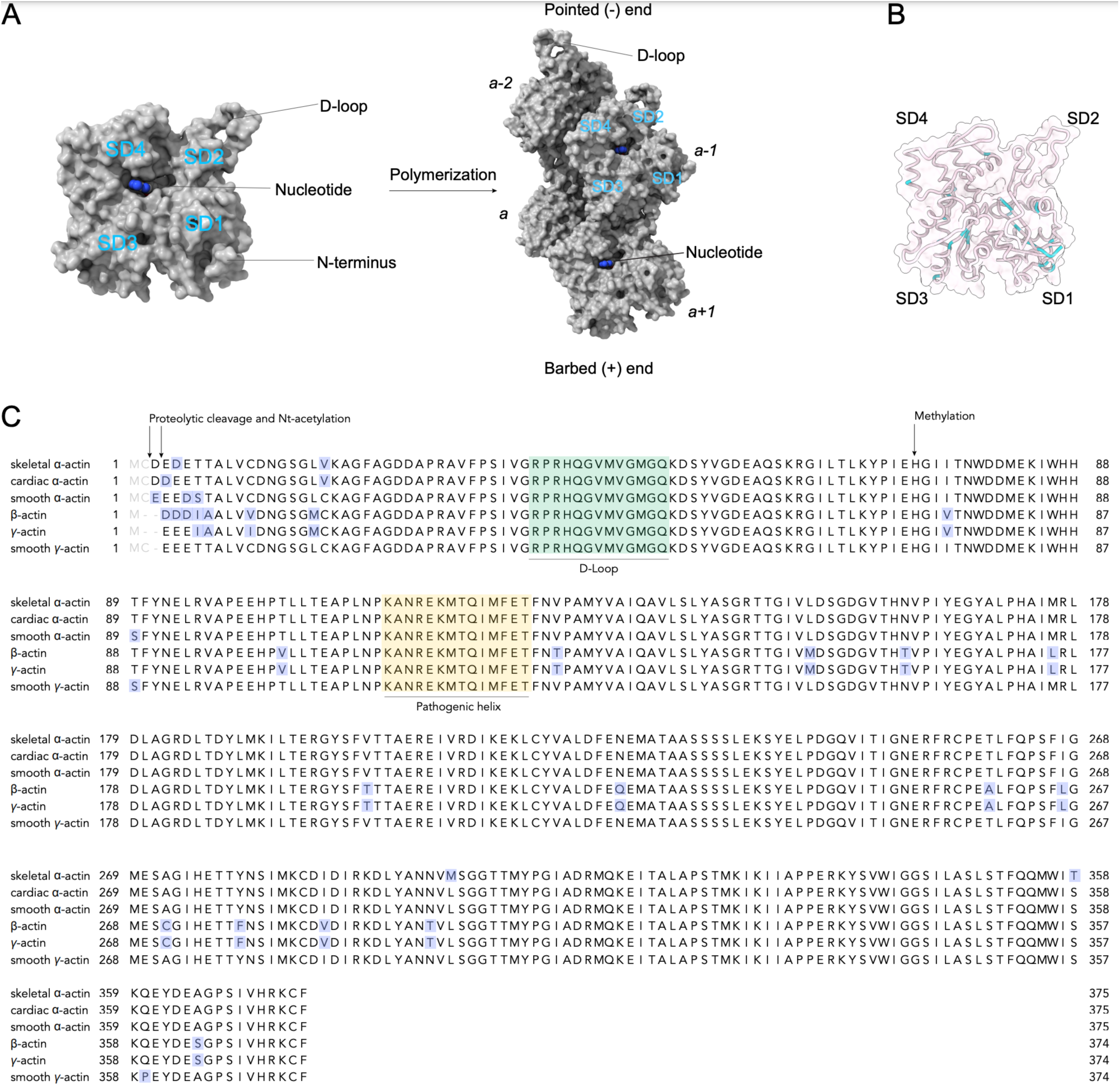
Sequence conservation in actin isoforms. (A) Classical view of the monomer structure of actin. Key regions are indicated. Monomers self-assemble into helical, polarized filaments. The arrangement of individual actin protomers in a filament are shown and the relative position denoted with italic numbers. The pointed (-) and barbed (+) ends are indicated. Monomer and filament structure not drawn to scale. (B) ConSurf analysis shows the conservation of amino acid positions in actin isoforms. Variations are shown in cyan color. (C) Sequence alignment of actin isoforms: rabbit skeletal α-actin (UniProt ID: P68135), bovine cardiac α-actin (UniProt ID: Q3ZC07), human smooth α-actin (UniProt ID: P62736), human β-actin (UniProt ID: P60709), human γ-actin (UniProt ID: P63261), human smooth γ-actin (UniProt ID: Q3ZC07). Conserved amino acids as shown in black, variable amino acids are highlighted in blue. Amino acids that are absent in mature actin isoforms are shown in grey. Key structural regions and PTMs visible in our cryo-EM structures are indicated.

**Figure 1-figure supplement 2:**
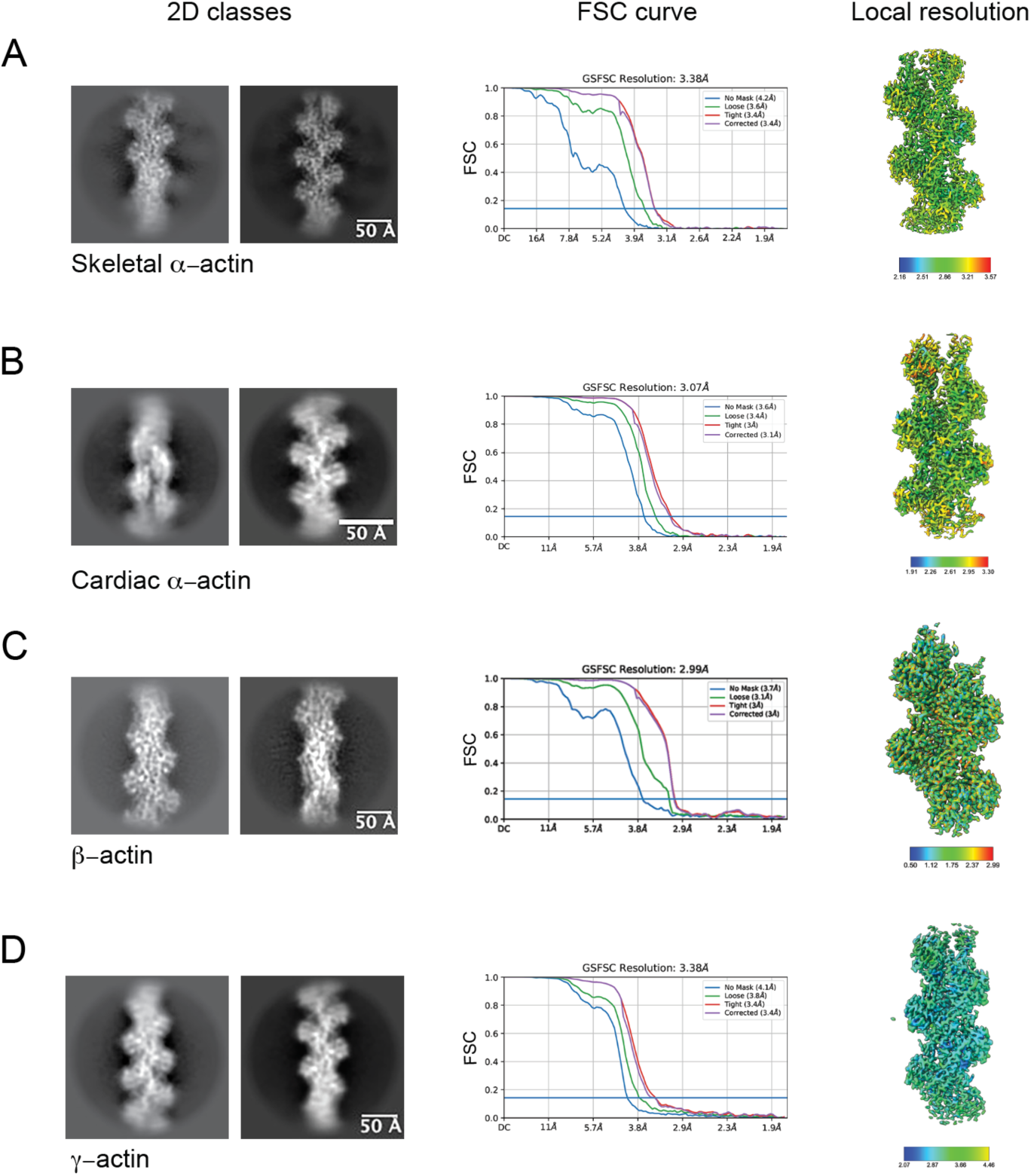
Image processing summary for actin isoforms. (A-D) 2D classes, global map-model FSC curves, and the local resolution estimated of the experimental maps are shown for skeletal α-actin (A), cardiac α-actin (B), β-actin (C), and γ-actin (D). The gold standard, FSC 0.143 criterion was used to estimate the global resolution of actin isoforms. The local resolution gradient is in Angstrom.

**Figure 3-figure supplement 1:**
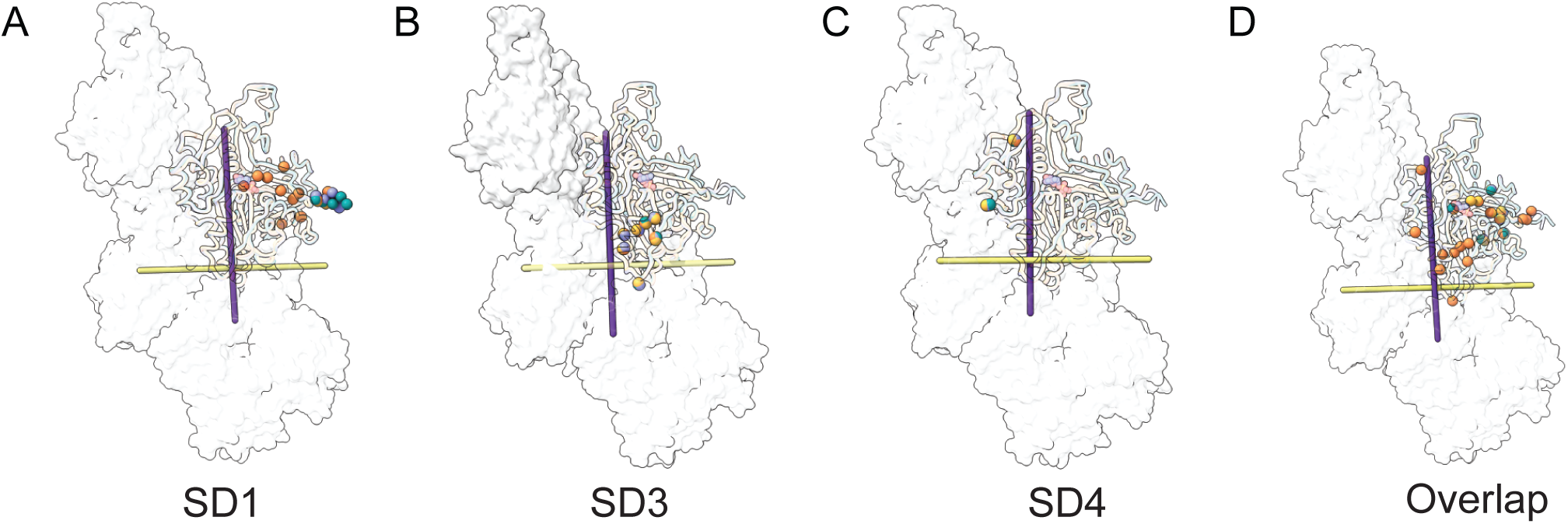
Variable amino acid distributions along the actin helical pitches. Non-conserved amino acids are shown along with the long pitch (purple axis) and short-pitch helix (yellow) for actin subdomains (A) SD1, (B) SD3, and (C) SD4. (D) Distribution of all variable amino acids in SD1, SD3, and SD4 for all actin isoforms.

